# Developing High-Yield and Safe Therapeutic EVs by Ablating Tissue Factor-Mediated Toxicity

**DOI:** 10.64898/2025.12.25.696171

**Authors:** Qingshan Luo, Xiang Li, Moxuan Yang, Xiaoyu Li, Shengya Xu, Tian Jiang, Junxian Xia, Jinfeng Ding, Xi Qin, Tong Zhao

## Abstract

The clinical translation of extracellular vesicles (EVs) as next-generation drug delivery vehicles is currently stalled by two interrelated challenges: the difficulty of manufacturing EVs at an industrial scale ("the yield problem") and the risks of toxicity associated with systemic administration of high doses ("the safety ceiling"). In this comprehensive study, we present a two-pronged genetic engineering strategy to overcome these limitations. First, we discovered that overexpression of glycosylphosphatidylinositol (GPI)-anchored proteins, such as CD55 and CD59, significantly enhances EV production in HEK293T and Expi293F cells. By engineering a truncated CD55 variant (TR3) that retains the GPI-anchor but lacks functional domains, we developed the ExoBoost cell line, which boosts EVs yield by approximately 50-fold without altering vesicle morphology. Second, we observed acute toxicity in mice following intravenous administration of high-dose ExoBoost EVs, with symptoms resembling venous thromboembolism. Through comparative analysis of EVs derived from other cell types, including keratinocyte stem cells (KSCs) and mesenchymal stem cells (MSCs), we identified Tissue Factor (TF/*F3*) as the critical causative agent of this lethal response. Moreover, we completely abolished this toxicity by generating a *F3*-knockout cell line (ExoBoost2.0) using CRISPR/Cas9, thereby raising the safe dose to as high as 1.5E12 particles per mouse, as quantified by nano flow-cytometry (nFCM). Finally, using a novel CD46-nanoluciferase reporter system, we demonstrated that high-dose administration of these safe EVs saturates hepatic and splenic clearance pathways, leading to a dramatic increase of systemic EVs accumulation, particularly in hard-to-target tissues such as the brain (∼10,000-fold) and muscle (∼1,000-fold). Additionally, repeated high-dose administration of ExoBoost2.0 EVs did not upregulate inflammatory cytokines or production of IgG and IgM. In summary, these findings establish a scalable, safe, and highly biocompatible EVs platform, potentially revolutionizing the drug delivery system in clinical application.

## Introduction

The advent of the genetic medicine era has generated an unprecedented demand for sophisticated delivery technologies. While adeno-associated viruses (AAVs) and synthetic lipid nanoparticles (LNPs) have demonstrated remarkable clinical success, their limitations are becoming increasingly apparent [^1–3^]. AAVs are restricted by a cargo capacity of approximately 4.7 kb and are associated with high immunogenicity, which typically precludes repeated administration. Furthermore, although AAV vectors are generally non-integrating, the risk of genomic insertion remains a concern. Conversely, while LNPs can efficiently encapsulate large nucleic acid sequences, they often exhibit limited ability to traverse biological barriers, possess poor protein packaging capacity, and suffer from immunogenic concerns associated with synthetic lipids. Against this backdrop, extracellular vesicles (EVs) have emerged as a promising alternative delivery platform, representing nature’s own intercellular communication system [^3^].

EVs are lipid-bilayer nanoscale vesicles secreted by nearly all cell types and can be classified as exosomes or ectosomes based on biogenesis [^4–6^]. The complex composition of EVs, featuring hundreds of naturally occurring surface proteins and lipids, facilitates self-recognition and potentially enhances bioavailability while minimizing immunogenicity. However, the clinical translation of EV therapeutics is hindered by two fundamental challenges: inefficient secretion, which prevents scalable production, and undefined safety profiles that restrict therapeutic dosing. To address the limitation of low productivity, various methods involving chemical regulation, physical stimulation, and physiological modification have been explored [^7–9^]. However, the mechanisms underlying EV secretion enhancement have not been thoroughly investigated; a systematic comparison of stimulants regarding downstream effects is necessary to identify a common mechanism and enable the creation of genetically enhanced EV secretion.

Additionally, while preliminary safety data exists [^10–20^], the toxicity profile of systemically administered EVs—particularly regarding the Maximum Tolerated Dose (MTD)—remains incompletely understood. This gap in knowledge is largely due to the lack of sufficient EV production required for rigorous high-dose toxicity testing. In the current limited-dose setting, less than 5% of injected EVs persist in circulation within minutes, resulting in short plasma half-lives [^21–23^]. They are primarily captured by the liver and spleen via the mononuclear phagocyte system (MPS), where the unique microarchitecture accelerates the clearance [^22–25^]. While inflammation can increase CNS uptake [^26,27^], under normal conditions less than 1% of injected EVs cross the blood–brain barrier [^28,29^]. Thus, increasing EV production and establishing a safe high - dose window to enhance delivery to target tissues beyond the liver and spleen, particularly across the blood-brain barrier, is a pivotal direction for EV-based therapeutics.

Here, we present the development and validation of the ExoBoost2.0 platform. By integrating a novel GPI-anchored protein-mediated EVs yield enhancement technology with the ablation of the *F3* (Tissue Factor) gene, this platform enables the industrial-scale production of non-thrombogenic EVs suitable for high-dose therapeutic applications and targeted delivery to challenging tissues such as the brain and muscle.

## Results

### Discovery of GPI-Anchored Proteins as Potent Enhancers of EV Biogenesis

Extracellular vesicles (EVs) carry conserved proteins, such as tetraspanins (CD81, CD63, CD9), TSG101, SDCBP, and ALIX, which mediate their core biological activities. They also contain cell-type-specific components that confer specialized functions [^30–34^]. Stem cells—including mesenchymal stem cells (MSCs) and keratinocyte stem cells (KSCs)—which possess intrinsic therapeutic properties, represent valuable sources of native EVs for research [^33,35^]. The HEK293 cell line, a human embryonic kidney derivative, has been widely employed in research and biopharmaceutical production for over three decades. Its advantages include rapid growth in chemically defined media, scalability, and high transfection efficiency, making it suitable for recombinant protein and therapeutic development [^36^]. Moreover, HEK293 cells are commonly used as a model system for generating EV-based drug delivery vectors due to their high EV productivity and immunologically neutral phenotype [^19,25^].

During a screen of engineered EV surface proteins in HEK293T cells, we serendipitously observed a substantial increase in EV production upon overexpression of the GPI-anchored proteins CD55 and CD59. These proteins, along with the single-pass transmembrane protein CD46, are key complement regulators [^37–40^]. Although all three are detected on EVs from diverse cell types, their roles in EV biogenesis or function remain unclear. We overexpressed each protein in HEK293T cells and isolated EVs from the conditioned medium. Western blot analysis of whole-cell lysates and purified EVs confirmed strong overexpression of each factor, along with the canonical EV markers ALIX and TSG101 (Figure S1A). GAPDH served as a loading control and, as expected, was markedly depleted in EV fractions. EV quantification by nano-flow cytometry (nFCM) was expressed as particles per liter of original conditioned medium. Cells overexpressing the GPI-anchored proteins CD55 and CD59 produced significantly more particles than wild-type (WT) HEK293T cells. In contrast, CD46 overexpression yielded EV numbers indistinguishable from WT cells (Figure S1B). This result strongly suggests that the enhanced EV biogenesis is not a general consequence of protein overexpression or complement regulation, but is specifically linked to the GPI-anchor moiety.

To test whether the GPI-anchor itself drives this effect, we engineered a series of truncated CD55 variants. Four constructs (TR1–TR4) were designed, each retaining the C-terminal signal sequence for GPI-anchoring but lacking progressively larger portions of the extracellular sushi domains (Figure S1C). All constructs, including full-length CD55, carried an N-terminal HA tag for uniform detection. These were transfected into Expi293F cells—a suspension-adapted, high-density culture derivative cell line of HEK293 optimized for protein production. Western blot analysis confirmed successful expression and efficient sorting of all truncated variants into EVs. Importantly, all GPI-anchored truncations, irrespective of extracellular domain size, significantly increased EV yield (Figure S1E). This finding highlights the potential of genetic engineering to enhance EV production and provides mechanistic insight into EV biogenesis. Based on its robust EV-promoting activity, we selected the TR3 variant for stable expression in Expi293F cells, thereby establishing the ExoBoost cell line for further development.

### Systemic Administration of High-Dose ExoBoost EVs Induces Acute Lethal Toxicity in C57BL/6 Mice

To ensure that the higher EV yield did not compromise vesicle quality, we performed a comprehensive comparison between ExoBoost EVs and Expi293F EVs. Nano-flow cytometry (nFCM) and nanoparticle tracking analysis (NTA) showed that comparable size distributions: peak diameters were 95.34 nm (ExoBoost EVs) vs 102.6 nm (Expi293F EVs) by nFCM, and 193.7 nm vs 153.3 nm by NTA (Figure 1A,B). Notably, size distributions and particle concentrations obtained by nFCM and NTA are known to vary due to differences in detection principles, sensitivity, and analytical limitations, as reported in prior methodological studies [^41,42^]. Unless otherwise specified, all particle concentrations reported in this study were determined using an Apogee Micro-GxP flow cytometer. To more accurately determine the size of ExoBoost EVs, we employed cryo-electron microscopy (cryo-EM), which allows visualization of EVs in a near-native, hydrated state. Cryo-EM confirmed the presence of intact lipid-bilayer vesicles, with statistical analysis revealing an average diameter of 102.67 nm (Figure S1F). Furthermore, western blot analysis verified the enrichment of canonical EV markers (ALIX, TSG101, CD9, CD81) and the absence of the mitochondrial contaminant CYC1 in both EV preparations (Figure 1C, 1D). Negative-stain transmission electron microscopy (TEM) confirmed typical cup-shaped, spherical membrane-bound morphology (Figure 1E, 1F).

**Figure 1.**
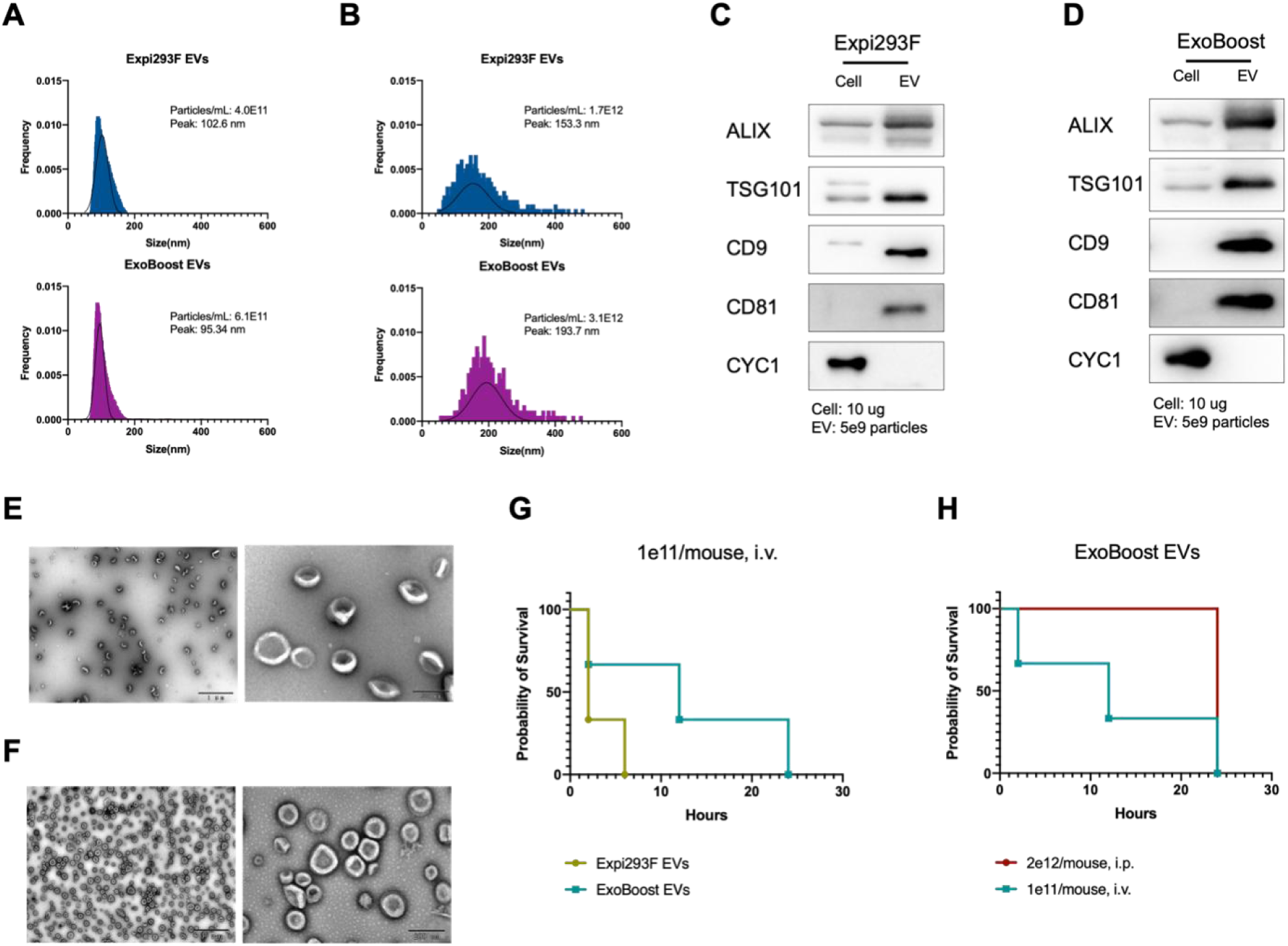
Characterization and *in vivo* toxicity of EVs isolated from Expi293F and ExoBoost cells. **(A)** Particle concentration and size distribution of Expi293F EVs, as determined by nano-flow cytometry (nFCM; upper panel) and nanoparticle tracking analysis (NTA; lower panel). **(B)** Particle concentration and size distribution of ExoBoost EVs, as determined by nFCM (upper panel) and NTA (lower panel). **(C)** Western blot analysis of EV biomarkers in cells and their corresponding EVs from Expi293F cells. **(D)** Western blot analysis of EV biomarkers in cells and their corresponding EVs from ExoBoost cells. **(E)** Transmission electron microscopy (TEM) images of Expi293F EVs, showing a large field view (left) and a zoomed-in view (right). **(F)** TEM images of ExoBoost EVs, showing a large field view (left) and a zoomed-in view (right). **(G)** Survival curve of C57BL/6 mice following intravenous (i.v.) injection of 1E11 particles of either Expi293F EVs or ExoBoost EVs (N=3). **(H)** Survival curve of C57BL/6 mice comparing the toxicity of ExoBoost EVs administered via intravenous (i.v.) injection of 1E11 particles versus intraperitoneal (i.p.) injection of 2E12 particles (N=3).

After overcoming production bottleneck with the ExoBoost platform, we utilized the improved yield to conduct high-dose *in vivo* safety studies. EVs were purified by density-gradient ultracentrifugation to reduce non - membrane particle contamination. Initially, C57BL/6 mice received a single intravenous (i.v.) tail-vein injection of 1E11 particles of either Expi293F or ExoBoost EVs. Strikingly, both groups exhibited an acute, severe toxic response: mice rapidly developed respiratory distress and ataxia, which progressed to convulsions and death within 30 minutes, resulting in 100% mortality (Figure 1G). The rapid and uniform onset of symptoms suggested a systemic vascular event rather than a delayed cellular toxicity. To test this hypothesis, we administered a separate cohort of mice with a substantially higher dose (2E12 particles per mouse, 20 times the lethal i.v. dose) via intraperitoneal (i.p.) injection. Remarkably, all animals receiving i.p. injection survived with no observable adverse effects (Figure 1H). This route allows slower absorption of EVs into the systemic circulation via the lymphatic system, avoiding the sudden peak plasma concentration achieved by i.v. injection. Together, these results strongly implicate a threshold-dependent, plasma-mediated mechanism as the cause of the acute lethal toxicity following intravenous administration.

### The Protein Corona of ExoBoost EVs Mediates the Observed Acute Toxicity

Proteins, particularly membrane-associated proteins, constitute a major component of EVs and are poised to mediate rapid biological responses through enzymatic or cascade reactions. Given the acute onset of toxicity following high-dose EV administration, we hypothesized that proteins on the EV surface (the protein corona) play a critical role in this process. To test this, we digested surface-exposed proteins on ExoBoost EVs with the broad-spectrum serine protease Proteinase K (ProK). Western blot showed complete loss of surface markers such as CD9 and the HA-tagged TR3 booster, while the luminal marker TSG101 remained intact, indicating that the lipid bilayer protected internal cargo from proteolysis (Figure 2A). nFCM analysis using an anti-HA-488 antibody confirmed a drastic reduction in HA signal on ProK-treated EVs (Figure 2B). Furthermore, TEM imaging confirmed that the "shaved" EVs retained their vesicular morphology and size distribution (Figure 2C).

**Figure 2.**
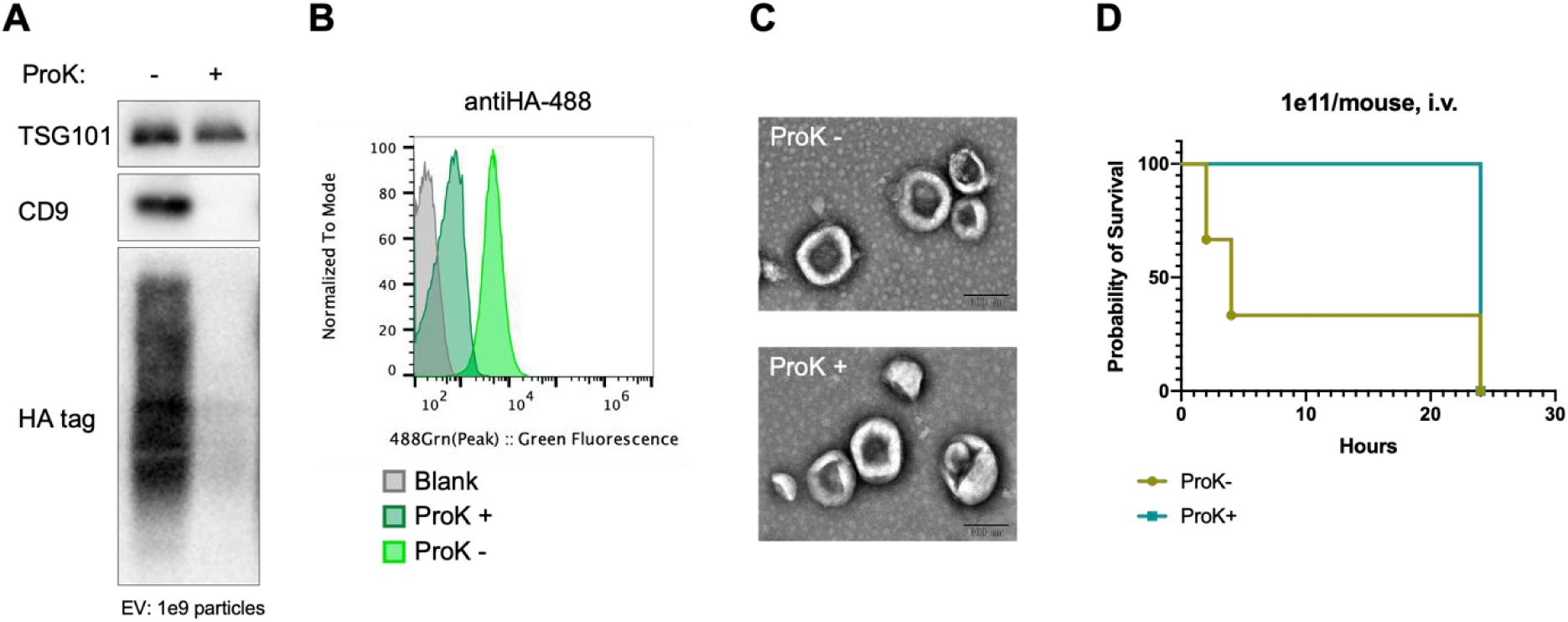
Characterization and in vivo toxicity of ExoBoost EVs with or without Proteinase K treatment. **(A)** Western blot analysis of EV biomarkers in ExoBoost EVs following treatment with or without Proteinase K (ProK). **(B)** Surface expression levels of the HA-tag on ExoBoost EVs with or without ProK treatment, as quantified by nFCM. **(C)** Morphology of ExoBoost EVs with or without ProK treatment, as visualized by TEM. **(D)** Survival curve of C57BL/6 mice following intravenous (i.v.) injection of 1E11 particles of ExoBoost EVs, with or without ProK pretreatment (N=3).

Crucially, when the surface protein digested ExoBoost EVs were administered intravenously at the previously lethal dose (1E11 particles per mouse), all animals survived with no observable adverse effects (Figure 2D). This result provides definitive evidence that the acute toxicity strictly dependent on surface proteins, rather than on lipids, internal RNA cargo, or buffer components.

### Tissue Factor is a Critical Mediator of ExoBoost EV Toxicity

The observed presentation—rapid respiratory distress and ataxia progressing to convulsions—closely resembled acute massive pulmonary thromboembolism [^43^]. This similarity led us to hypothesize that the EVs could act as systemic pro-coagulant agents. Coagulation *in vivo* is primarily initiated via the extrinsic pathway, triggered by Tissue Factor (TF) [^44–47^]. Under physiological conditions, TF is sequestered in the subendothelial layer and not exposed to circulating blood. Direct exposure of TF to plasma Factor VIIa rapidly activates the coagulation cascade, potentially causing disseminated intravascular coagulation. Notably, TF is a single-pass transmembrane protein previously reported to be present on EVs [^48,49^].

To investigate the potential role of TF in ExoBoost EV toxicity, we first quantified TF expression in ExoBoost cells and their EVs. Keratinocyte stem cells (KSCs) and their EVs served as a positive control, as KSCs are known to express high levels of TF—a physiological adaptation for rapid hemostasis during wound healing [^50^]. As predicted, KSCs expressed higher baseline levels of TF than ExoBoost cells. More importantly, TF was highly enriched in EVs derived from both KSCs and ExoBoost cells (Figure 3D), suggesting that KSC EVs might induce toxicity at a significantly lower dose. Subsequent *in vivo* dose-response toxicity assays confirmed this prediction. While ExoBoost EVs were non-toxic at a dose of 5E10 particles per mouse, KSC-EVs caused 100% mortality at doses as low as 2.5E9 particles per mouse (Figure 3E, 3F). This approximately 100-fold increase in toxic potency correlates directly with the higher TF content carried by KSC-EVs, strongly implicating TF as a key driver of the acute coagulopathic toxicity.

**Figure 3.**
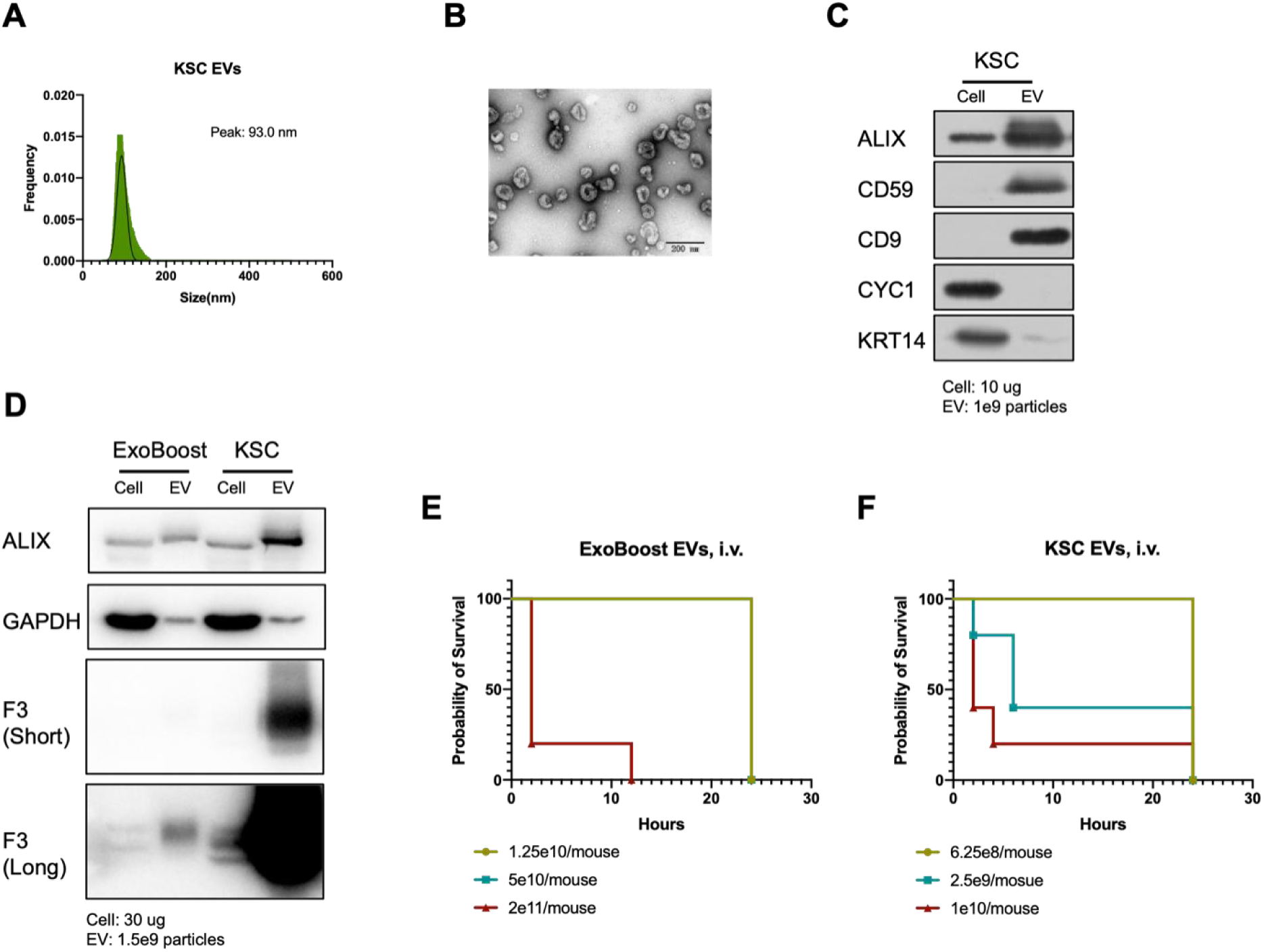
Tissue Factor (TF) is a key mediator of EV-induced toxicity in mice. **(A)** Size distribution of KSC EVs as determined by nFCM. **(B)** Morphology of KSC EVs visualized by TEM. **(C)** Western blot analysis of EV biomarkers in KSCs and their derived EVs. **(D)** Western blot analysis of TF expression levels of ExoBoost EVs and KSC EVs. **(E)** Survival curve of C57BL/6 mice following intravenous (i.v.) injection of ExoBoost EVs at doses of 1.25E10, 5E10, or 2E11 particles per mouse (N=5). **(F)** Survival curve of C57BL/6 mice following intravenous (i.v.) injection of KSC EVs at doses of 6.25E8, 2.5E9, or 1E10 particles per mouse (N=5).

### ExoBoost2.0: F3 Knockout Ablates Toxicity While Maintaining High EV Yield

Having identified TF as the key mediator of toxicity, we engineered a next-generation producer cell line designed to retain the high-yield phenotype of the ExoBoost platform while eliminating TF expression. Using CRISPR/Cas9, we knocked out TF in Expi293F cells to generate a clonal *F3*-knockout (F3KO, clone #12) line, confirmed by the absence of TF protein (Figure S2A-D).

The TR3 booster construct was then stably expressed in this F3KO clone to restore high EV productivity, creating the cell line designated ExoBoost2.0. Flow cytometry confirmed robust TR3 expression in ExoBoost2.0 cells, which maintained the ∼50-fold yield increase over the original Expi293F line, with total output and yield per cell comparable to the original ExoBoost cells (Figure S2E, S2F).

ExoBoost2.0 EVs exhibited a similar size distribution (peak diameter ∼96.9 nm by nFCM) and morphology compared to ExoBoost EVs (Figure 4A, 4B). Western blot analysis confirmed the complete absence of TF in ExoBoost2.0 EVs (Figure 4C). For *in vivo* testing, we encountered a practical limitation: the physical solubility of highly concentrated EV preparations. EV solutions became opalescent at ∼1E11 particles/mL and viscous at ∼1E13 particles/mL (Figure 4D), making it difficult to administer more than 2×10¹² particles in a standard 200 µL intravenous bolus. This suggests a physical solubility limit may be reached before the biological toxicity threshold.

**Figure 4.**
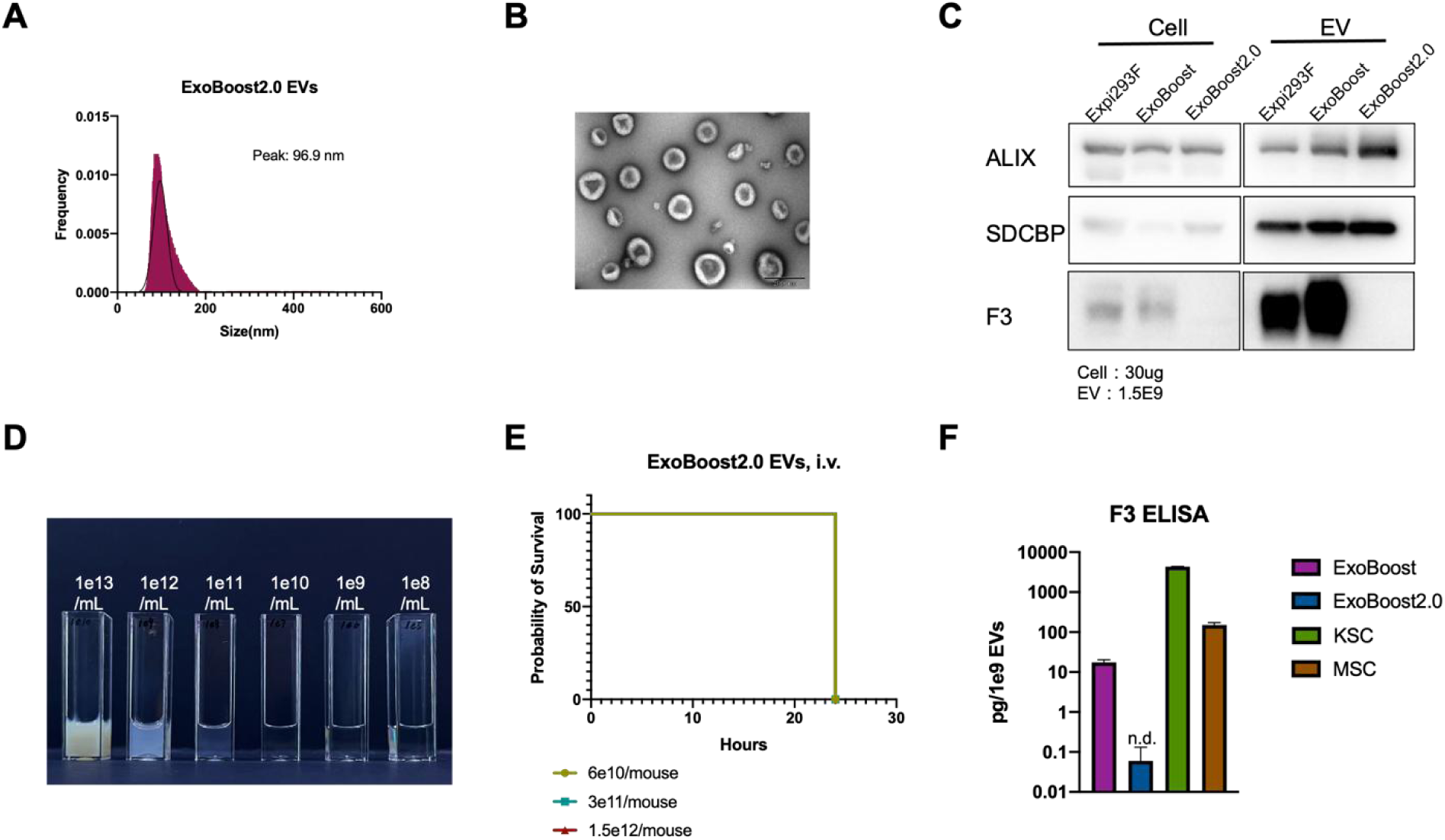
Tissue Factor knockout abolishes acute toxicity of high-dose intravenous injection of EVs. **(A)** Particle size distribution of ExoBoost2.0 EVs as determined by nFCM. **(B)** Morphology of ExoBoost2.0 EVs visualized by TEM. **(C)** Western blot analysis of TF and EV biomarkers in cells and corresponding EVs from Expi293F, ExoBoost, and ExoBoost2.0 cell lines. **(D)** Physical appearance of concentrated ExoBoost2.0 EV preparations, appearing milky white at 1E11 particles/mL and slightly viscous at 1E13 particles/mL. **(E)** Survival curve of C57BL/6 mice following intravenous (i.v.) injection of ExoBoost2.0 EVs at doses of 6×E10, 3E11, or 1.5E12 particles per mouse (N=5). **(F)** Quantification of Tissue Factor protein levels in 1E9 particles of EVs derived from ExoBoost, ExoBoost2.0, KSCs, and MSCs, as measured by ELISA.

Despite this, dose-escalation toxicity studies in C57BL/6 mice showed that intravenous administration of ExoBoost2.0 EVs, at doses ranging from 6×E10 to 1.5E12 particles per mouse, induced no observable adverse effects—even at the highest dose, which is 15-fold higher than the lethal dose (1E11 particles/mouse) of the original ExoBoost EVs (Figure 4E). By ablating TF expression, we thus increased the maximum tolerated dose (MTD) by at least 30-fold, effectively removing a major biological safety barrier for the systemic administration of high-dose EVs.

We further confirmed the presence of TF in EVs derived from mesenchymal stem cells (MSCs), which have been associated with thrombotic risks in clinical reports [^51^]. MSCs were expanded in chemically defined medium and characterized by standard surface markers within early passages (P8) to minimize serum-derived artifacts (Figure S3A). EVs were isolated and characterized by nFCM and TEM (Figure S3B, S3C). Western blot analysis confirmed that MSC EVs carry substantially higher levels of TF than Expi293F EVs, consistent with the high endogenous TF expression in MSCs (Figure S3D), indicating a potential toxicity risk at high doses of MSC EVs injection.

To establish a standardized, quantifiable metric for comparison across cell sources, we quantified the absolute amount of TF protein (ng) per 1E9 EV particles using ELISA for EVs derived from ExoBoost, ExoBoost2.0, KSCs, and MSCs. This TF content index provides a critical parameter for manufacturing quality control and safety assessment. Beyond protein quantity, evaluating TF coagulation activity is essential and constitutes part of our ongoing work. While genetic editing in primary cells like KSCs and MSCs remains challenging, alternative strategies—such as small-molecule inhibitors or using differentiated iPSC-derived progenitors—could be employed to reduce TF levels in their EVs, enhancing safety profiles for therapeutic applications.

### Biodistribution and Pharmacokinetics of ExoBoost2.0 EVs Following High-Dose Systemic Administration

The safety of high dose EV administration enabled us to investigate whether saturating the body’s natural clearance pathways could enhance EVs delivery to secondary tissues. To track EVs *in vivo* without the artifacts of lipophilic dyes, we developed a reporter system based on Nanoluc (nLuc) fused to the transmembrane and membrane-proximal domains of CD46 at its C-terminus, ensuring luminal encapsulation (Figure S4A). The reporter construct was then overexpressed in both ExoBoost and ExoBoost2.0 cells. EVs from these lines (ExoBoost_nLuc and ExoBoost2.0_nLuc) were characterized by nFCM and TEM, confirming expected size and morphology (Figure S4B, S4C). Western blot analysis verified successful nLuc expression in cells and its enrichment in corresponding EVs (Figure S4D). Importantly, Proteinase K treatment did not diminish luciferase activity, confirming efficient luminal encapsulation (Figure S4E). The bioluminescence signal demonstrated excellent linearity with particle number across a 6-log range, validating the reporter for quantitative studies (Figure S4F).

Using this reliable system, we mapped the biodistribution and pharmacokinetics (PK) of ExoBoost2.0 EVs at various doses. First, a low dose (2.5E10 particles/mouse) of ExoBoost2.0_nLuc or ExoBoost_nLuc EVs was administered intravenously. Tissues harvested one hour post-injection (with saline perfusion to remove blood-pool signal) showed predominant EVs accumulation in the liver and spleen, consistent with known clearance by the MPS [^52,53^] (Figure 5A). The plasma concentration and tissue signals for ExoBoost2.0_nLuc EVs were moderately higher, likely due to the absence of coagulation-mediated rapid clearance, leading to prolonged circulation. We then performed a dose-escalation study, administering ExoBoost2.0_nLuc EVs at doses ranging from 2.5E10 to 1E12 particles/mouse. At the highest dose, total tissue luminescence increased by approximately 1,000-fold, despite only a 40-fold increase in particle number (Figure 5B). Notably, EV accumulation in the brain increased by ∼700-fold with 10-fold dose increased by comparing the 1E12 and 1E11 dose groups. This disproportionate enhancement supports that a massive bolus temporarily overwhelms the clearance capacity of the liver and spleen, extending the circulation half-life and enabling greater partitioning into tissues with less permeable endothelium, such as the brain and muscle.

**Figure 5.**
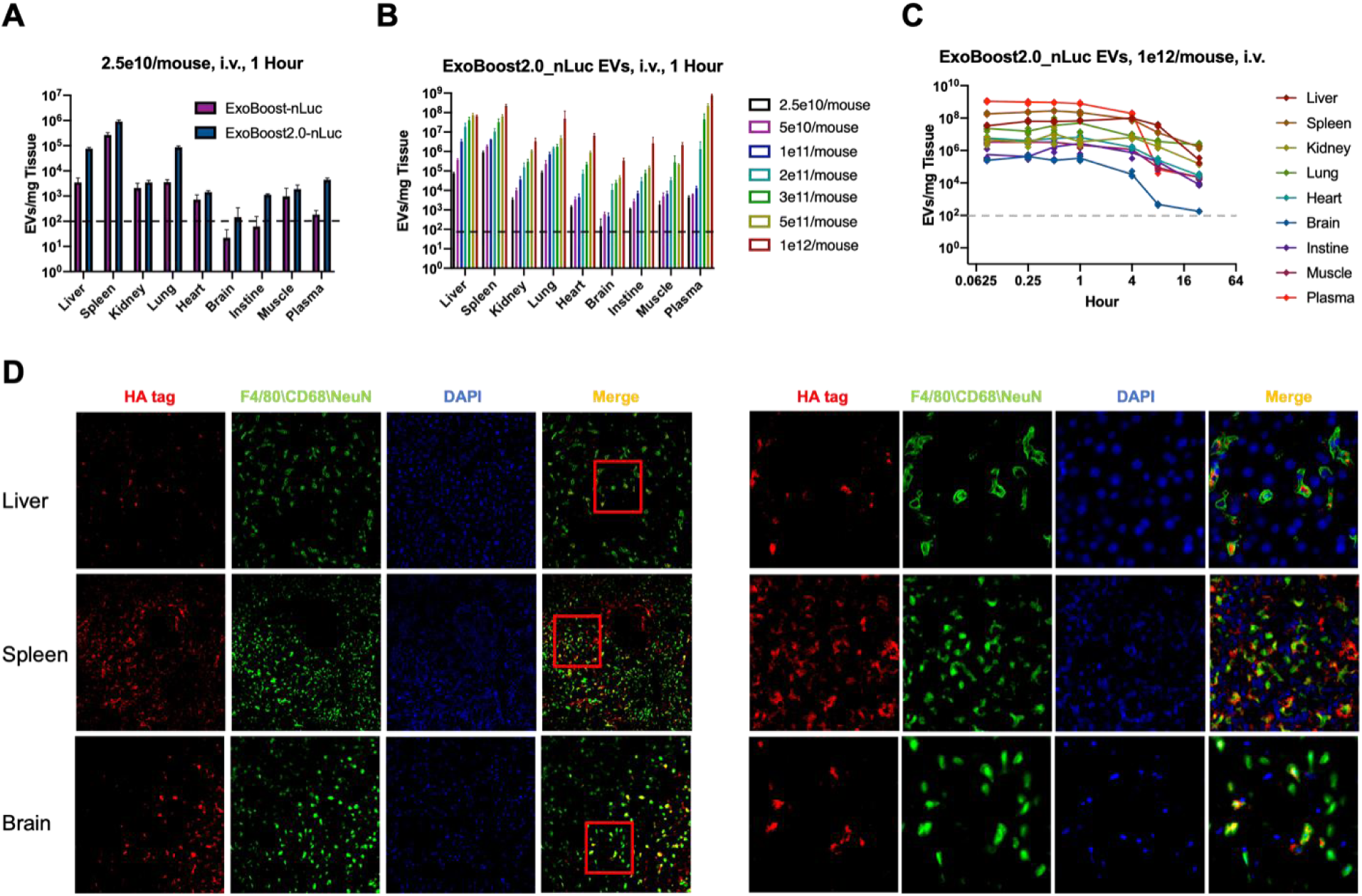
*In vivo* pharmacokinetics and biodistribution of ExoBoost and ExoBoost2.0 EVs carrying the nano-luciferase reporter. **(A)** Whole-body biodistribution of ExoBoost_nLuc and ExoBoost2.0_nLuc EVs at 1-hour post-intravenous (i.v.) injection of 2.5E10 particles in C57BL/6 mice (N=3 for ExoBoost_nLuc, N=4 for ExoBoost2.0_nLuc). **(B)** Biodistribution of ExoBoost2.0_nLuc EVs at 1-hour post-i.v. injection at varying doses, ranging from 2.5E10 to 1E12 particles per mouse (N=4). **(C)** Pharmacokinetics of ExoBoost2.0_nLuc EVs in organs over 24 hours following i.v. injection of 1E12 particles per mouse (N=4). **(D)** Representative immunofluorescence images showing the cellular localization of HA-tagged ExoBoost2.0_nLuc EVs (red) within the liver (co-stained with F4/80 for macrophages), spleen (co-stained with CD68 for macrophages), and brain (co-stained with NeuN for neurons), nuclei are counterstained with DAPI (blue), showing a large field view (left) and a zoomed-in view (right).

Next, a 24-hour PK study using the high dose (1E12 particles/mouse) showed sustained EV signals in tissues over the entire period, indicating successful tissue retention rather than transient passage (Figure 5C). Pharmacokinetic analysis comparing low-dose ExoBoost_nLuc EVs with high-dose ExoBoost2.0_nLuc EVs revealed a remarkable increase in the area under the curve (AUC), particularly in the brain, where AUC was enhanced by up to 30,000-fold (Figure S5). This dramatic increase highlights the potential of high-dose EV administration for enhancing drug delivery to the brain.

To visualize intra-organ distribution, we performed immunofluorescence staining of tissue sections using an anti-HA antibody (targeting the TR3 booster) alongside cell-specific markers. EV signal largely co-localized with F4/80⁺ (Kupffer cells in the liver) and CD68⁺ (Macrophages in the spleen) cells, confirming MPS-mediated clearance (Figure 5D). High-resolution imaging showed EV signals as punctate structures within endosomal compartments. Notably, EV signals were also detected beyond macrophages, particularly in the spleen, warranting further investigation into alternative recipient cells and functional consequences. Most significantly, we observed HA-positive puncta co-localizing with NeuN⁺ neurons in the cortex, providing direct evidence that at high doses, EVs can cross the blood-brain barrier and deliver cargo to neuronal cells.

### Minimal Immunogenicity Upon Repeated High-Dose Administration of ExoBoost2.0 EVs

Following the demonstration of acute safety, we evaluated potential immunogenicity after repeated exposure. Mice received six weekly intravenous injections of 1E12 particles of ExoBoost2.0 EVs (Figure 6A). ELISA of plasma after the final injection showed no significant elevation in total mouse IgG or IgM compared to PBS-treated controls (Figure 6B). While some studies report EV-specific antibody responses without a rise in total immunoglobulin [^25^], our initial data indicate no massive humoral immune activation. Furthermore, quantitative PCR analysis of key inflammatory cytokines (*Il6, Il1b, Il10*) in the liver, spleen, and kidney showed no upregulation compared to PBS controls (Figure 6C). Together, these results confirm that repeated high-dose administration of ExoBoost2.0 EVs does not elicit significant antibody production or systemic inflammatory cytokine activation.

**Figure 6.**
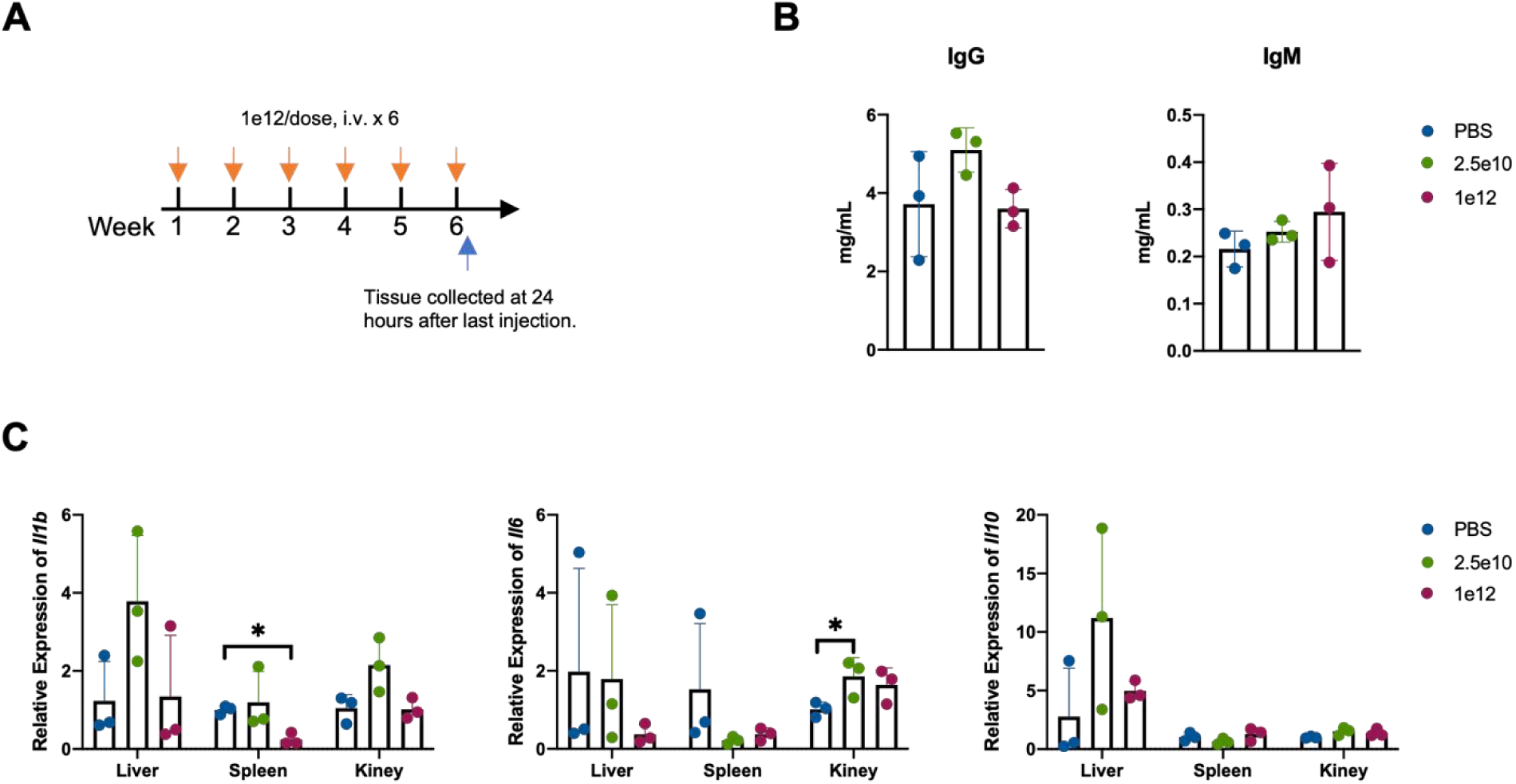
Evaluation of inflammatory response and immunogenicity following multi-dose intravenous administration of ExoBoost2.0 EVs in mice. **(A)** Schematic diagram of the multiple-dosing study. **(B)** Plasma concentrations of anti-EV immunoglobulin G (IgG) and immunoglobulin M (IgM) antibodies, as measured by ELISA (N=3). **(C)** mRNA expression levels of inflammatory cytokines in the liver, spleen, and kidney, as determined by quantitative real-time PCR (N=3).

## Discussion

The development of the ExoBoost2.0 platform directly addresses two of the most pervasive challenges in extracellular vesicle therapeutics: manufacturing scalability and systemic administration safety. By elucidating and manipulating the fundamental biology underlying vesicle biogenesis and coagulation, we have engineered a system capable of delivering therapeutic quantities of EVs to difficult-to-access tissues. Our discovery that GPI-anchored proteins potently drive EV biogenesis provides a novel, genetically simple handle for bioprocess engineering. GPI-anchors preferentially partition into the liquid-ordered (Lₒ) phase of the plasma membrane (lipid rafts). We posit that the high-density clustering of our overexpressed TR3 protein induces local membrane curvature or facilitates EVs secretion. This mechanism is elegant in its simplicity, requiring only the introduction of a short GPI-anchoring signal sequence—a feature that renders this yield-enhancing strategy maybe highly transferable across diverse cell types.

The identification of Tissue Factor (TF) as the principal mediator of acute EV toxicity resolves a critical gap in EV research. This lethal effect has likely been underreported due to the low doses typically administered in preclinical studies. Our findings carry profound implications for therapies based on MSC EVs, which are currently the most advanced in clinical translation. Our data indicate that MSCs produce EVs with significant TF content—a level that may be safe at low doses but poses a serious thrombotic risk when doses are escalated to those required for robust organ targeting. Furthermore, donor-to-donor variability in TF expression could underlie the inconsistent safety profiles sometimes observed in clinical settings. We therefore propose that TF quantification become mandatory quality control (QC) metrics for all clinical-grade EV batches. Besides, our gene-editing strategy offers a clear roadmap for creating "universally safe" therapeutic cell sources, such as iPSC-derived MSCs or KSCs, where the *F3* gene is ablated at the pluripotent stage prior to differentiation—an approach we are actively pursuing.

The central nervous system represents one of the most fortified therapeutic frontiers. While strategies employing specific targeting ligands (e.g., Rabies Virus Glycoprotein) have shown promise in enhancing brain delivery, absolute efficiency remains low. Our study introduces a paradigm-shifting alternative: sheer mass action. By eliminating the primary toxicity barrier, we can administer previously impossible doses (e.g., 1E12 particles/mouse). This high-dose regimen transiently saturates the scavenging capacity of the liver and spleen. Once these primary clearance "sinks" are overwhelmed, the systemic exposure (AUC) of circulating EVs increases dramatically, thereby statistically enhancing their probability of interacting with and traversing the brain endothelium. The orders-of-magnitude increase in brain accumulation underscores that achieving high-dose safety is, in itself, a powerful passive targeting strategy. Coupling this safe, high-yield ExoBoost2.0 platform with active targeting moieties (e.g., peptides, ligands, aptamers) presents a highly synergistic path forward for CNS drug delivery.

While ExoBoost2.0 constitutes a significant advance, several challenges remain on the path to clinical translation. A universal hurdle for all nano-delivery systems, including EVs, LNPs, and viral vectors, is the efficient endosomal escape and cytosolic release of therapeutic cargo (e.g., siRNA, mRNA, proteins). Future work must therefore focus not only on loading these abundant, safe vesicles with diverse therapeutics but also on engineering them to overcome intracellular trafficking barriers. Additionally, further refinement of targeting specificity and a deeper understanding of the long-term fate and potential immunogenicity of repeatedly administered engineered EVs are essential. In summary, by solving the foundational problems of scalable production and acute toxicity, the ExoBoost2.0 platform transforms the EV therapeutic landscape, turning the previous bottleneck of mass production into a new engine for targeted delivery.

## Methods and Materials

### Cell Culture

293T cells and its derivatives were cultured in DMEM supplemented with 10% of fetal bovine serum and maintained in a humid incubator with 5% CO2 at 37 ℃. For EVs isolation, fetal bovine serum was replaced with EV-deleted serum by ultracentrifuge at 150, 000 xg for 18 hours. Expi293F cells and other derivatives such as ExoBoost and ExoBoost2.0 cells, were cultured in serum-free medium in the Erlenmeyer flasks shaking at 120 rpm in a humid incubator with 5% CO2 at 37℃. Keratinocyte stem cells were cultured on tissue-culture treated flasks in serum-free EpiZero medium. Mesenchymal stem cells were cultured in medium supplemented with 5% serum replacement for expansion and cultured in chemical defined medium for EVs isolation. No antibiotics are used and cells are routinely tested for mycoplasma negative to ensure the reliable culture system.

### Plasmid Construction and Stable Cells Selection

To generate a stable cell line overexpressing the polypeptide of interest, PippyBac (PB) transposon system or pLenti lentivirus system was used. Briefly, a nucleic acid fragment encoding a coding sequences (CDS) of the polypeptide of interest was synthesized (Genewiz) and cloned into expression vectors. For HEK293T cells, approximately 5E5 of cells were transfected 2 µg of PB Transposon vector carrying the nucleic acid fragments of the polypeptide in combination with 1 µg of PB Transposase vector by using Lipofectamine 3000, according to the manufacturer’s instructions. The transfected cells were then incubated and selected under 200 µg /mL of hygromycin B for 10 days to obtain a stable cell line overexpressing the gene of interest. For lentivirus packaging, pLenti, pM2G, pVSVG were co-transfected to HEK293T cells using Lipofiter and culture for 48 hours before lentivirus harvest. Lentivirus were isolated by co-precipitation with reagent. Lentivirus containing pellet was resuspended in PBS and store at - 80 ℃. To infection with targeted cells, each lentivirus were added to cells with appropriated MOI for 48 hours, then cells were selected under 100 ug/mL hygromycin B or 10ug/mL puromycin for 10 days to ensure the stable expression. Before cell banking, RCL was tested as negative by qPCR method and p24 ELISA.

### EVs Isolation

For volume less than 2L, cell culture supernatant was harvested by centrifugation with 500 x g for 10 min at 4℃ to remove the liable cells. Then supernatant was transferred and subjected to the centrifugation with 2500 x g for 20 min at 4℃ to remove the dead cells. Then the supernatant was filtered through a 0.22-μ m membrane to remove cell debris as well as big extracellular vesicles. Then EVs will be pelleted by the ultracentrifugation at 100,000 x g for 85 min at 4℃. The pellet was pooled and washed with PBS followed by a second ultracentrifugation at 100,000 x g for 85 min at 4℃. Ultimately, the supernatant was discarded and the pellet was resuspended in PBS followed with the final purification by size exclusion chromatography (SEC) column.

For volume more than 2L, culture supernatant was harvested by centrifugation with 500 x g for 10 min at 4℃ to remove the liable cells. Then supernatant was transferred and subjected to the centrifugation with 2500 x g for 20 min at 4 ℃ to remove the dead cells. Then the supernatant was filtered through a 0.45/0.22-μm membrane to remove cell debris as well as big extracellular vesicles. Then the clarified supernatant was concentrated by tangential flow filtration using mPES hollow fibre. After Benzonase treatment, crude EVs were further concentrated with hollow fibre again followed with purification by SEC. Ultracentrifugation was performed if high-concentration EVs required.

For highly purified EVs preparations, crude pellets were diluted to 3 mL total volume with PBS, mixed with 9 mL of 60% OptiPrep, and added into the bottom of a 38-mL tube. Lower-density solutions were prepared by diluting with homogenization buffer (250 mM sucrose, 10 mM Tris-HCl, 1 mM EDTA, pH 7.4) to yield final iodixanol concentrations (vol/vol) of 30%, 23%, and 18%. Successive layers 9 mL of 30%, 6 mL of 23%, and 6mL of 18% iodixanol solutions were added on top carefully. Finally, 3 mL of PBS was added to the top of the gradient. The gradient preparation was ultracentrifuged for 16 h with 150,000x g at 4C. The EVs layer was carefully extracted and concentrated by ultracentrifugation. The EV pellet was resuspended in PBS and aliquoted at ∼1E10 particles/mL, then stored at −80 C.

### Western Blot

Cells and EVs were lysed in lysis buffer. Protein concentration was measured using a BCA protein assay kit for cells to ensure equal loading, and EVs were loaded with equal particles. The samples were resolved by SDS-PAGE, followed by transferring onto a PVDF membrane. Membranes were probed with respective primary antibodies (Table S1). Bound primary antibodies were recognized by HRP-linked secondary antibodies (CST). Immunoreactivity was detected by ECL Star. Digital images of films were taken with ChampChemi 610 (Sagecreation).

### Nanoparticle tracking analysis

EVs particle size and concentration were assessed by Brownian motion with the ZetaView (Particle Metrix) based on manufacturer’s protocol. Samples were diluted at room temperature in 3 mL PBS within the operational range and injected into the instrument. The motion of particles was monitored, and recorded videos were analyzed with ZetaView software. Raw .fcs document was analyzed with FlowJo and Prsim software for statistical analysis.

### Nano-Flow Cytometry analysis

EVs particle size and concentration were detected on an Apogee MicroGxP Flow Cytometer that was specially developed for the analysis of nanoparticles. In some cases, before or during the sample measurements, two commercially available mixes of beads were used as controls and calibration standards (Apogee 1493 and Apogee 1517, Apogee Flow Systems, UK). EV samples were diluted in 1 mL PBS within the operational range and subjected to the instrument. Raw .fcs document was analyzed with FlowJo and Prsim software for statistical analysis.

### Transmission Electron Microscopy (TEM) and Cryogenic Electron Microscopy (Cryo-EM)

For negative staining TEM, EVs were diluted in PBS to an optimal concentration. A 5µL aliquot was deposited onto 230-mesh carbon-coated copper grids. After incubation for 1 min at room temperature, excess liquid was removed using filter paper. The grids were then negatively stained with 1.5% uranyl acetate or 1% phosphotungstic acid for 1 min. After air-drying, the samples were imaged using a JEOL JEM-1200EX transmission electron microscope operating at an acceleration voltage of 100 kV.

For Cryo-EM analysis, 3.5 µL of the EVs was applied to Quantifoil R1.2/1.3 holey carbon grids. The grids were blotted for 3.5 s at 100% humidity and 4 °C using a FEI Vitrobot Mark IV, and immediately plunged into liquid ethane for vitrification. The vitrified grids were imaged using a FEI Talos F200C transmission electron microscope.

### In vivo Assay

Male C57BL/6J mice (8 – 10 weeks old) were purchased from Beijing Vital River Laboratory Animal Technology Co., Ltd. (Beijing, China). All animals were housed in a specific pathogen-free (SPF) facility under a standard 12-hour light/12-hour dark cycle with ad libitum access to food and water. All animal experiments were approved by the Animal Care and Use Committee of TheraXyte Bioscience.

To evaluate the toxicity of EVs, mice received a tail - vein injection of either low - dose or high - dose suspended in 200 µL PBS. Mice response to EVs within 30min was recorded with videos for dose-dependent tests. Mice status were examined in 2, 6, 12, 24 hours after injection and survival curves were analyzed by Prism.

To evaluate the biodistribution and PK of EVs, mice received a tail - vein injection of either low - dose or high - dose suspended in 200 µL PBS. At different time points post-injection, mice were anesthetized with 1.5% tribromoethanol (20 µL g⁻¹ body weight). Blood (∼0.2mL) was collected from retro-orbital sinus into tubes containing heparin sodium to prevent clotting. Plasma was isolated by centrifugation at 1,000× g for 10 min. After blood collection, mice were perfused transcardially with ice - cold PBS to flush residual blood from the circulation. Major organs and tissues were excised, weighed, and placed in lysis buffer with zirconia beads. Tissues were homogenized using a bead - beating homogenizer. Nanoluc luciferase activity in both plasma and tissue homogenates was quantified with the Nano-Glo® Luciferase Assay System (Promega, USA) according to the manufacturer’s protocol. The bioluminescence intensity was used to quantify the EV content in each sample.

### Immunofluorescence staining of tissues

Post-injection, mice were perfused, and the harvested organs were immediately fixed in tissue fixative, then processed and paraffin - embedded. Liver/spleen (transverse) and brain (coronal, ∼Bregma −2.30 mm) sections were deparaffinized and blocked with 3% BSA. Sequential immunofluorescence was performed via iterative cycles of antibody incubation and elution. For each target, sections were incubated with primary antibodies (overnight, 4 °C). To track EVs, an anti-HA tag antibody was used. Cell types were labeled using Anti-F4/80, Anti-CD68, and Anti-NeuN. Following PBS washes, HRP-conjugated secondary antibodies were applied, and signals were enhanced using a TSA reaction (10 min, RT). Between staining cycles, antibody complexes were eluted to prevent cross-reactivity. Finally, sections were stained with DAPI, treated with an autofluorescence quencher, and mounted. Imaging was performed on a NIKON ECLIPSE C1 system.

### Immunogenicity Assessment

Total plasma IgG and IgM levels were quantified using a Mouse IgG ELISA Kit and a Mouse IgM ELISA Kit, respectively, following the manufacturer’ s instructions. Briefly, plasma samples and standards were added to plates pre-coated with capture antibodies and incubated for 2 hour at 25 °C with orbital shaking at 200 rpm. Following incubation, a detection antibody was added for another hour. A chromogenic substrate was then applied, and the absorbance was measured using a Synergy HTX multi-mode reader.

Cytokine gene expression was analysis by RT-qPCR. Total RNA was extracted from tissue homogenates using the TRIzol reagent. The RNA was reverse-transcribed into cDNA using supermix. Quantitative real-time PCR (qPCR) was performed to measure the mRNA levels of inflammatory cytokines, including Tnfa, Il6, Il1b, and Il10. Reactions were carried out using Hieff® qPCR SYBR Green Master Mix on an Applied Biosystems 7500 Fast Real-Time PCR System. Relative gene expression was calculated using the ΔΔCt method, normalized to GAPDH. The primer sequences are listed in Table S2.

### Statistical Analysis

Data are presented as Mean ± SD. Differences between groups were analyzed by T-test. For significance analysis, * represents for p-value <0.05 and ** represents for p-value < 0.01. Particle concentrations were taken as reported by the respective instruments, while the particle size distribution was obtained by re-synchronization of the raw data, and the peak values were derived from a Gaussian curve-fitting procedure.

## Author contributions

Conceptualization: T.Z.; Methodology: Q.L., X.L., M.Y. and T.Z.; Investigation: Q.L., X.L., M.Y., X.Y.L., S.X., T.J., J.X, J.D. and T.Z.; Resources: X.Y.L., J.X. and J.D.; Data Curation: Q.L., X.L., M.Y., S.X., T.J., and T.Z.; Writing - Original Draft: Q.L., M.Y., S.X., T.J. and T.Z.; Writing - Review & Editing: Q.L., X.L., M.Y., X.Q. and T.Z.; Visualization: Q.L., M.Y., X.Y.L., S.X., T.J., and T.Z.; Supervision: X.Q. and T.Z.

