## Supplementary Information for "Developing High-Yield and Safe Therapeutic EVs by Ablating Tissue Factor-Mediated Toxicity"

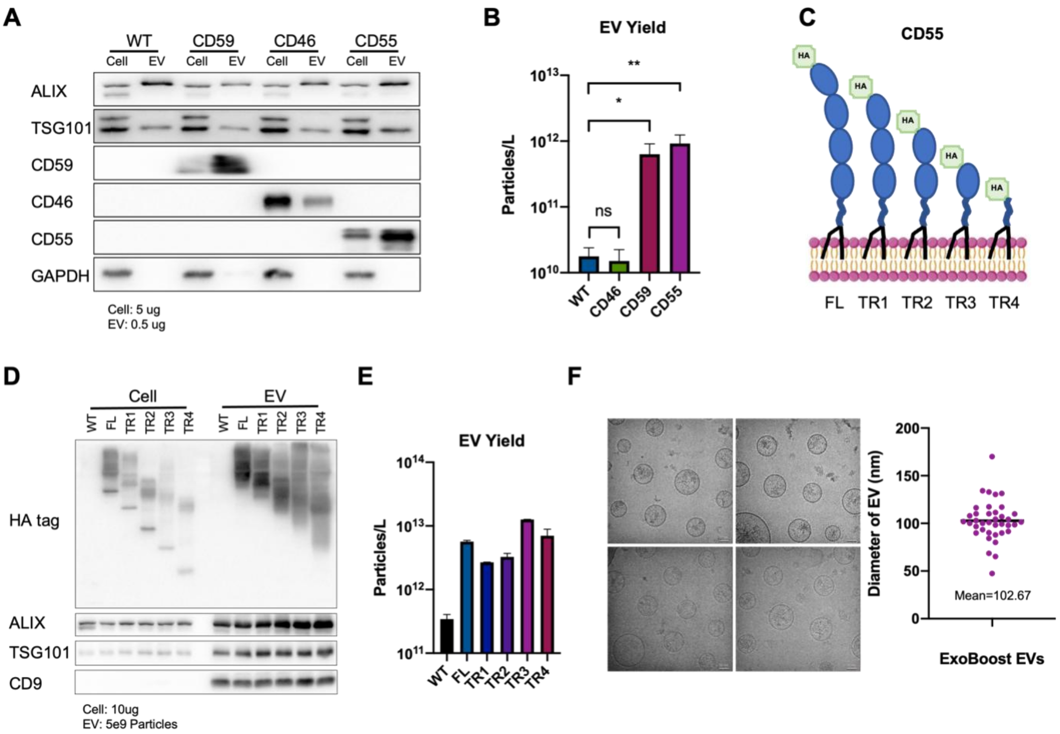


**Figure S1. Enhanced EV production in 293T and Expi293F cells via overexpression of GPI-anchored proteins CD55 and CD59. (A)** Western blot analysis of cells and corresponding EVs of wild-type (WT) 293T cells and 293T cells overexpressing CD59, CD46, or CD55. **(B)** Quantification of EV yield from WT 293T cells and 293T cells overexpressing CD46, CD55, or CD59, culture for 5 days. **(C)** Schematic diagram of full-length CD55 and truncated proteins TR1/2/3/4, HA-tag is fused to the N-terminal. **(D)** Western blot analysis of cells and corresponding EVs of WT Expi293F cells and Expi293F cells overexpressing full-length CD55 and truncated proteins TR1/2/3/4. **(E)** Quantification of EV yield from WT Expi293F cells and Expi293F cells overexpressing full-length CD55 and truncated proteins TR1/2/3/4, culture for 5 days. **(F)** Size distribution of ExoBoost (Expi293F cell overexpressing TR3) EVs, as determined by Cryo-electron microscopy (Cryo-EM) (N=38).

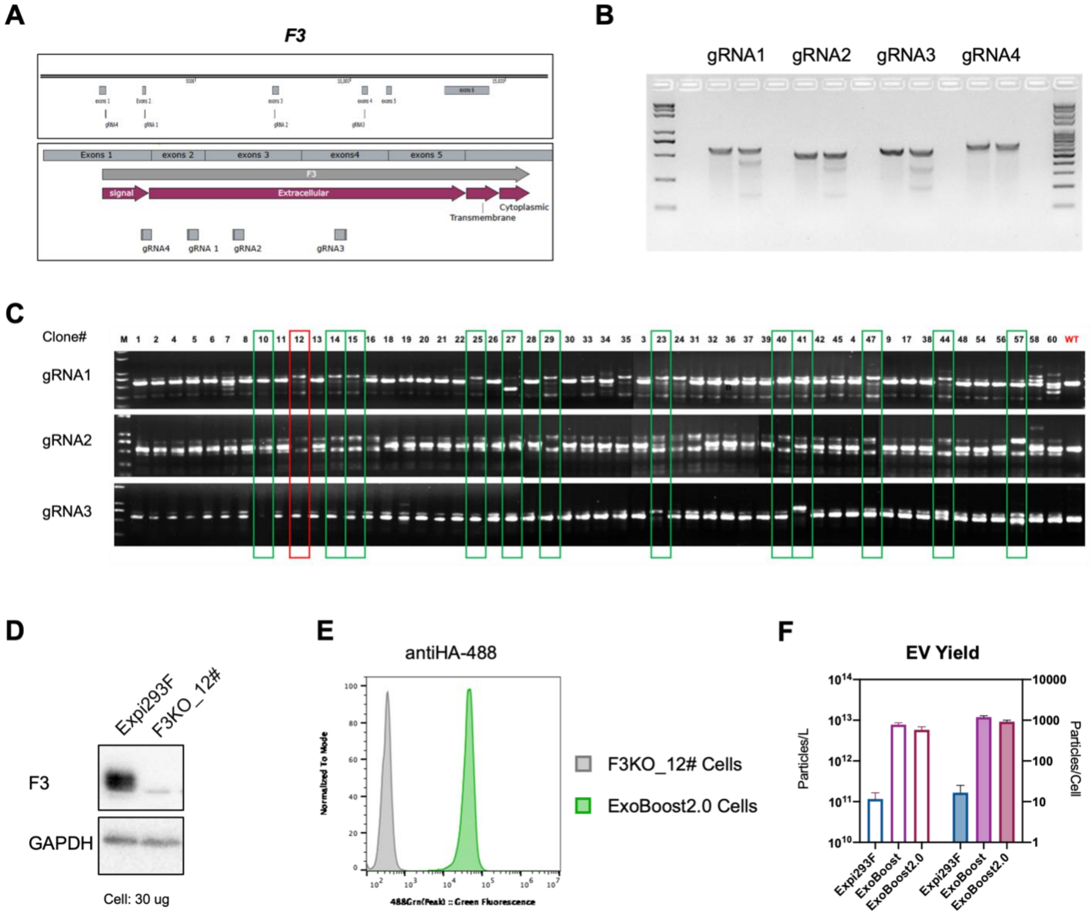


**Figure S2. Generation of Tissue Factor knockout Expi293F cells and construction of the ExoBoost2.0 cell line.** **(A)** Schematic of the human *F3* gene (encoding Tissue Factor) transcripts and the targeted loci of guide RNAs (gRNA 1-4). **(B)** Assessment of genome editing efficiency for individual gRNAs (gRNA 1-4) using a T7 endonuclease I (T7E1) cleavage assay. **(C)** Genotyping analysis of single-cell colonies derived from Expi293F cells co-transfected with gRNAs 1, 2, and 3. **(D)** Western blot confirmation of Tissue Factor completely knockout at the protein level in the selected clone #12. **(E)** Flow cytometry analysis confirm TR3 expression levels in the ExoBoost2.0 cells. **(F)** Quantification of EV yield from Expi293F, ExoBoost, ExoBoost2.0 cells, culture for 4 days.

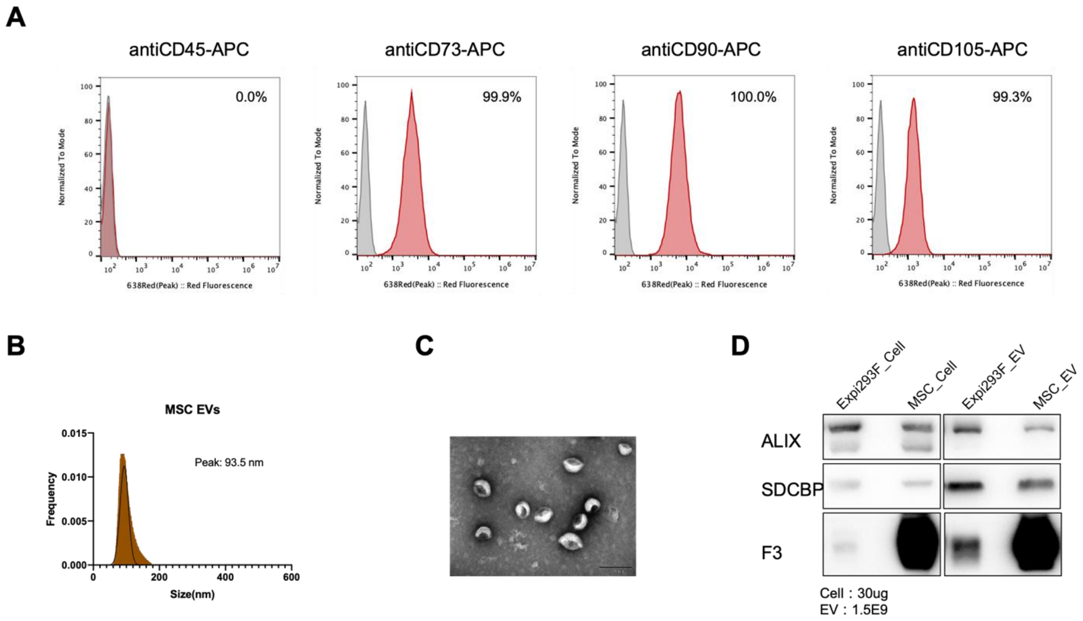


**Figure S3. Characterization of mesenchymal stem cells (MSCs) and their derived EVs. (A)** Flow cytometry analysis of standard MSC surface markers. **(B)** Size distribution of MSC-derived EVs, determined by nFCM. **(C)** Representative TEM image illustrating the morphology of MSC-derived EVs. **(D)** Western blot analysis of EV biomarkers and TF protein levels in MSCs and derived EVs, comparing to Expi293F cells and EVs.

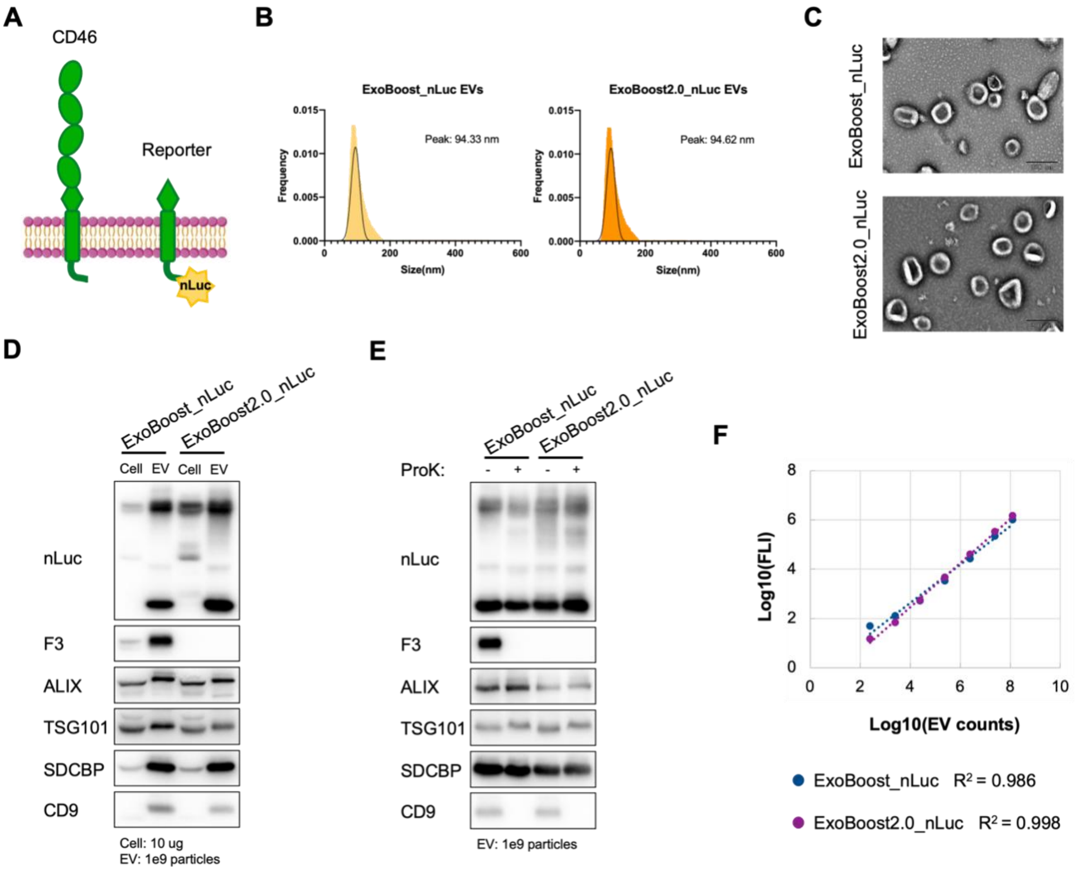


**Figure S4. Endogenous loading of nano-luciferase (nLuc) into EVs via a CD46-based scaffold in ExoBoost and ExoBoost2.0 cells. (A)** Schematic of the CD46-nLuc fusion reporter construct. The nano-luciferase is fused to the C-terminus of truncated CD46, ensuring its enrichment within the lumen of secreted EVs. **(B)** Size distribution of EVs derived from ExoBoost_nLuc and ExoBoost2.0_nLuc cells, as determined by nFCM. **(C)** Morphology of ExoBoost_nLuc and ExoBoost2.0_nLuc EVs, visualized by TEM. **(D)** Western blot analysis of nano-luciferase and EV biomarkers expression in cells and corresponding EVs from ExoBoost_nLuc and ExoBoost2.0_nLuc cell lines. **(E)** Western blot analysis of nano-luciferase and EV biomarkers in ExoBoost2.0_nLuc EVs following treatment with or without Proteinase K, demonstrating the luminal localization of nano-luciferase. **(F)** A linear correlation between the number of ExoBoost_nLuc and ExoBoost2.0_nLuc EVs and the luminescence intensity, validated over a range from 1E2 to 1E8 particles.

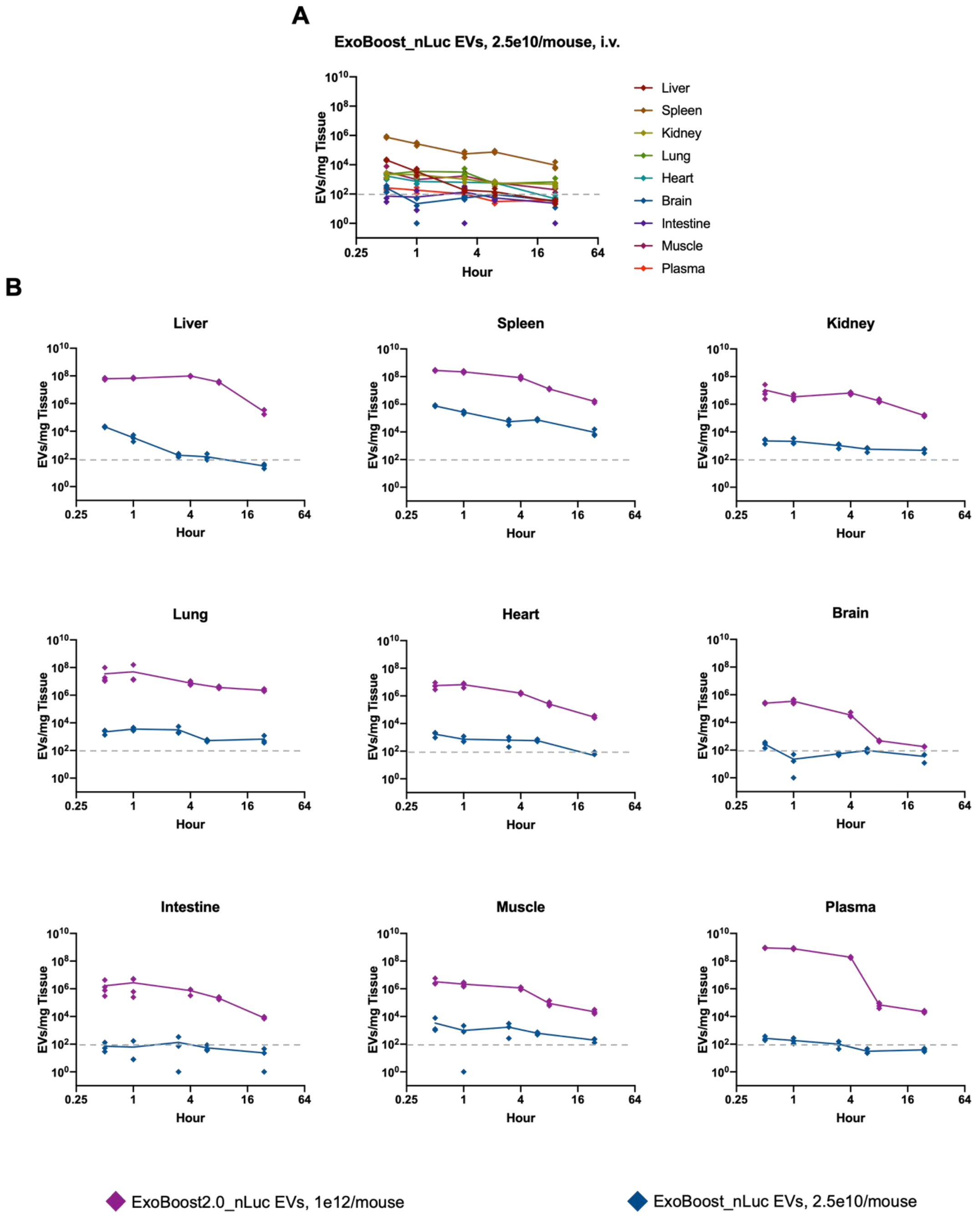


**Figure S5. Pharmacokinetics of ExoBoost_nLuc and ExoBoost2.0_nLuc EVs in individual tissues over 24 hours. (A)** Pharmacokinetics of ExoBoost_nLuc EVs in organs over 24 hours following i.v. injection of 2.5E10 particles per mouse. **(B)** Tissue distribution and clearance kinetics were quantified following a single intravenous (i.v.) injection of 2.5E10 particles of ExoBoost_nLuc EVs (blue) or 1E12 particles of ExoBoost2.0_nLuc EVs (purple) per mouse (N=3 for ExoBoost_nLuc,

N=4 for ExoBoost2.0_nLuc). Data for each organ are plotted separately to illustrate tissue-specific pharmacokinetic profiles.

**Table S1**. Antibody information

| No. | Antibody Name | Brand | Art.No. |
|---|---|---|---|
| **1** | Alix | abcam | ab186429 |
| **2** | TSG101 | abcam | ab125011 |
| **3** | CD46 | Santa Cruz | SC166159 |
| **4** | CD55 | Santa Cruz | SC51733 |
| **5** | CD59 | Santa Cruz | SC133170 |
| **6** | GAPDH | CST | 2118S |
| **7** | CD81 | abcam | ab79559 |
| **8** | NanoLuc | R&D | MAB10026 |
| **9** | HA tag | abcam | ab9110 |
| **10** | CD9 | abcam | ab236630 |
| **11** | CYC 1 | Proteintech | 10242-1-AP |
| **12** | TF | Santa Cruz | sc37441 |
| **13** | SDCBP | abcam | ab133267 |
| **14** | KRT14 | abcam | ab7800 |

**Table S2.** qPCR primer

| id | sequence |
|---|---|
| mIL-1βF | CAACCAACAAGTGATATTCTCCATG |
| mIL-1βR | GATCCACACTCTCCAGCTGCA |
| mGAPDHF | AACTTTGGCATTGTGGAAGG |
| mGAPDHR | ACACATTGGGGGTAGGAACA |
| mTNF-α-F | CATCTTCTCAAAATTCGAGTGACAA |
| mTNF-α-R | TGGGAGTAGACAAGGTACAACCC |
| mIL6-F | ACCAGAGGAAATTTTCAATAGGC |
| mIL6-R | TGATGCACTTGCAGAAAACA |
| mIL10-F | CGGGAAGACAATAACTGCACCC |
| mIL10-R | CGGTTAGCAGTATGTTGTCCAGC |
